# Coordinated topoisomerase function shapes the fluoroquinolone response of *Chlamydia trachomatis*

**DOI:** 10.64898/2026.03.19.712910

**Authors:** Andrew Terrebonne, Caitlynn Diggs, Scot P. Ouellette, Leiqiong Gao, Yuk-Ching Tse-Dinh, Li Shen

**Affiliations:** Department of Microbiology, Immunology, and Parasitology, Louisiana State University Health Sciences Center, New Orleans, LA 70112; Department of Pathology, Microbiology, and Immunology, University of Nebraska Medical Center, Omaha, Nebraska, USA; Biomolecular Sciences Institute, Department of Chemistry and Biochemistry, Florida International University, Miami, FL 33199, USA

**Author notes:** Equal contribution (Andrew Terrebonne, Caitlynn Diggs). Correspondence to: Li Shen, MD, Ph. D., Department of Microbiology, Immunology and Parasitology, Louisiana State University Health Sciences Center, New Orleans, LA 70112.

**Keywords:** DNA gyrase, TopA, DNA supercoiling, *Chlamydia trachomatis*, fluoroquinolone moxifloxacin, persistent form, DNA replication, transcription, antibiotic tolerance, and stress adaptation

## Abstract

DNA supercoiling is essential for the developmental cycle of *Chlamydia trachomatis*, yet its role in shaping antibiotic responses remains poorly understood. We investigated how the fluoroquinolone moxifloxacin (Mox) influenced *C. trachomatis* growth across developmental stages with its distinct supercoiling levels. Early Mox exposure completely halted bacterial growth, whereas treatment during mid-cycle produced enlarged, persistent forms and abolished formation of infectious progeny. These stage-specific outcomes coincided with inhibition of DNA replication, depletion of DNA gyrase, and transcriptional repression of *ompA* and *omcB*, accompanied by preserved or elevated expression of the stress-responsive *groESL1* operon. Mox also elicited compensatory downregulation of topoisomerase I (TopA), consistent with attempts to rebalance intracellular supercoiling. Together, these data demonstrate that fluoroquinolone susceptibility in *C. trachomatis* reflects stage-dependent supercoiling levels. Perturbation of supercoiling homeostasis drives developmental arrest and persistence phenotypes, highlighting coordinated gyrase-TopA activity as a key determinant of fluoroquinolone tolerance and a potential target for overcoming persistent infection.

**Importance:** *C. trachomatis*, a medically significant bacterial pathogen, can persist under antimicrobial pressure, complicating treatment strategy. This study links supercoiling homeostasis to fluoroquinolone tolerance, offering mechanistic insights into chlamydial adaptation to antibiotic stress and identifying potential targets to overcome persistence—an urgent challenge in global reproductive health.

## Introduction

*Chlamydia trachomatis* (Ctr) is a Gram-negative, obligate intracellular pathogen and the leading cause of bacterial sexually transmitted infections worldwide (1, 2). Although many infections are asymptomatic, untreated or chronic Ctr infection can lead to severe sequelae, including pelvic inflammatory disease, ectopic pregnancy, infertility, epididymitis, and reactive arthritis. Neonates born to mothers with untreated Ctr genital infections can develop severe conjunctivitis and pneumonia. Epidemiological studies further link Ctr infection to increased susceptibility to other sexually transmitted pathogens and to long-term complications such as cervical and ovarian cancers (3, 4). Despite the availability of effective antibiotics, persistent or recurrent infections remain a clinical challenge (5–7), and the underlying mechanisms that enable Ctr to survive antimicrobial pressure are not fully understood.

Ctr pathogenesis is closely tied to its unique biphasic developmental cycle (8, 9). Following entry into host mucosal epithelial cells, the infectious elementary body (EB) quickly decondenses its chromatin and differentiates into a noninfectious yet metabolically active reticulate body (RB) in a membrane-bound inclusion. RBs subsequently go into an exponential growth phase, or mid-cycle, from approximately 8-16 hours post-infection (hpi), before asynchronously undergoing secondary differentiation to EBs. EBs are ultimately released to infect new host cells. Under stress conditions (10–15), including nutrient deprivation, interferon-γ exposure, and certain antibiotic treatments, Ctr can enter a persistent state, which is characterized by aberrant RB-like persistent forms that remain viable but fail to produce infectious progeny. Persistence is reversible upon stress removal and is increasingly recognized as a contributor to chronic infection and treatment failure (16–20).

Progression through the Ctr developmental cycle relies on timed waves of transcription modulated by sigma factors (21–24), transcriptional regulators (25–29), and DNA topology (30–32). It is known that DNA supercoiling is primarily maintained by the opposing activities of topoisomerase I (TopA) and DNA gyrase in bacteria (33–36). Fluoroquinolones are broad-spectrum antibiotics that treat infections by targeting DNA gyrase and Topo IV. However, their broad use has been followed by emergence of resistance (37–41). Fluoroquinolone-resistant Ctr clinical isolates are rarely reported but are beginning to appear (18, 42). Experimental work indicates that exposure of Ctr to fluoroquinolones can select for resistance-associated mutations (43, 44) or induce Ctr to become metabolically dormant or persistent (5, 45).

Recent findings highlight the importance of balanced gyrase and TopA activity for Ctr development and antibiotic response (46, 47). Indeed, CRISPR interference (CRISPRi)-mediated repression of *topA* disrupts supercoiling homeostasis, arrests development, and sensitizes Ctr to fluoroquinolones. However, how lethal fluoroquinolone exposure perturbs DNA supercoiling and topoisomerase expression during infection, and how these changes shape Ct adaptation, remains unresolved.

In this study, we examine how the fluoroquinolone moxifloxacin (Mox) affects Ctr pathobiology across key developmental stages that differ in their intrinsic supercoiling levels (31–33). We characterize Mox-induced changes in bacterial growth, morphology, DNA replication, topoisomerase expression, and promoter-specific transcription in selective virulence genes. Collectively, these analyses define how disruption of DNA supercoiling dynamics influences the capacity of Ctr to adapt to topological stress and reveal coordinated topoisomerase regulation as a critical determinant of fluoroquinolone tolerance and persistence induction.

## Results

### Moxifloxacin exerts developmental stage–dependent effects on Ctr

To assess how the Ctr developmental stage influences the susceptibility of the pathogen to Mox, HeLa 229 cells (HeLa hereafter) were infected with strain L2/P_*ompA*_-GFP (48) and exposed to increasing drug concentrations at 0, 8, 16, or 24 hpi). These time points correspond to distinct DNA supercoiling states in Ctr: low supercoiling levels shortly after infection, increasing supercoiling during mid-cycle replication of RBs, peak supercoiling ~24 hpi, and declining levels during the late stage (30, 32).

Mox exposure at 0 or 8 hpi completely prevented inclusion formation, indicating high sensitivity during the early stage of the developmental cycle. Treatment at 16 hpi produced smaller inclusions containing enlarged, aberrant forms, accompanied by sharply reduced infectious EB progeny and lack of detection of the EB-associated protein OmcB (Figure 1a–d). In contrast, the addition of Mox at 24 hpi caused only mild defects.

**Figure 1.**
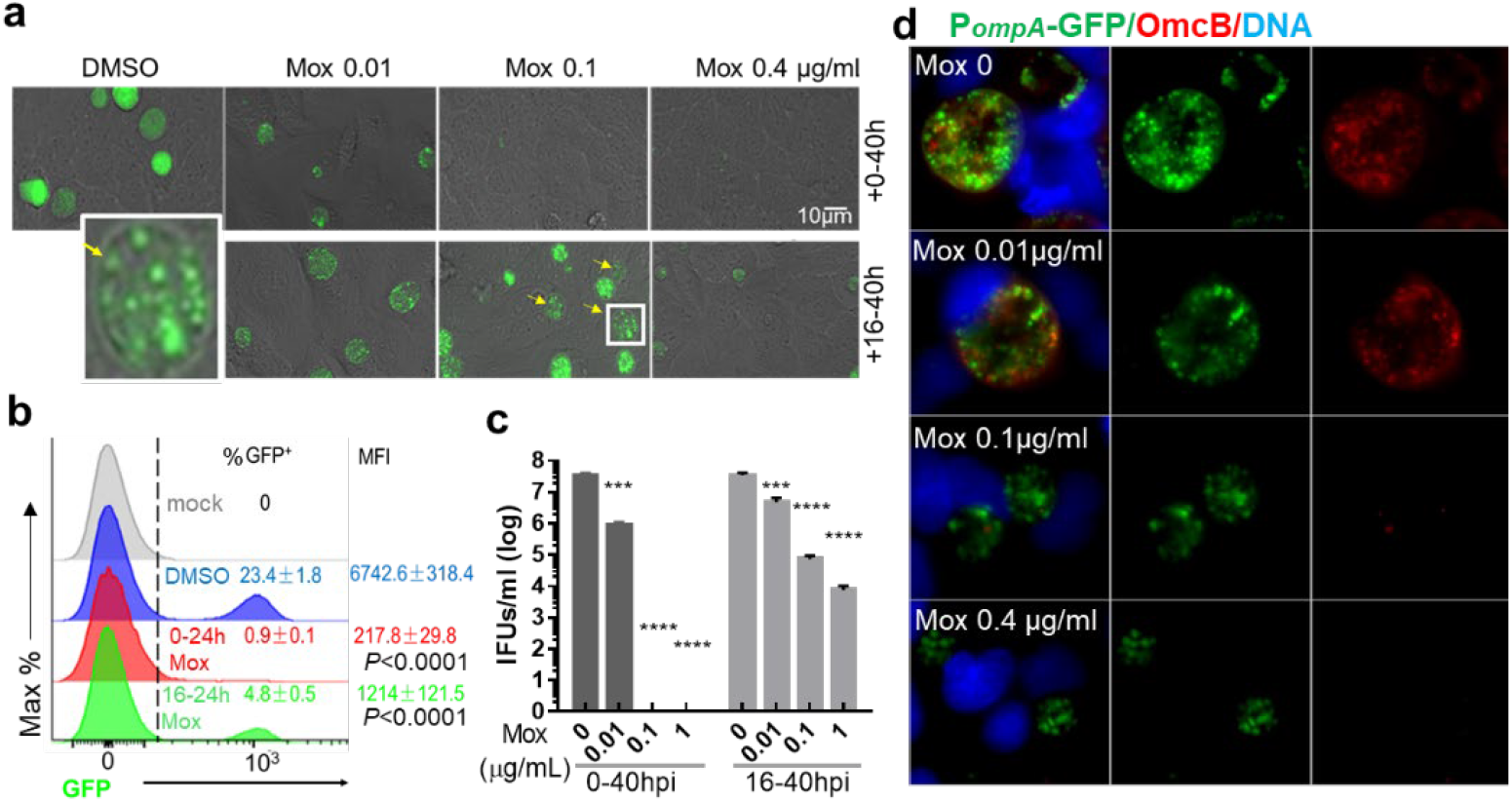
Time- and dose-dependent effects of Mox on *C. trachomatis* growth. HeLa cells were infected with L2/P_*ompA*_-GFP (48) at a multiplicity yielding ~30% infected cells, followed by treatment with increasing concentrations of Mox as indicated. (**a**) Live-cell fluorescence microscopy of chlamydial inclusions in cells with/without Mox for 40 h (0–40 hpi) or 24 h (16–40 hpi). Arrows indicate inclusions with aberrant persistent forms; inset shows magnified view. Scale bar, 10 µm. (**b**) Flow-cytometric quantification of GFP^+^ cells (%) and mean fluorescence intensity (MFI) after exposure to Mox at 0.1 µg/mL for 24 h (0–24 hpi) or 8 h (16–24 hpi). Mock-infected cells and DMSO-treated infected cells were used as controls. Data represent means ± standard derivation (SD) from three biological replicates, each measured in technical triplicate. Statistical comparison of MFI between exposed samples to DMSO control was performed using *t*-test, P<0.0001. (**c**) Enumeration of infectious EBs by inclusion-forming unit (IFU) assay. The IFU values represent means ± SD from three replicates in an experiment. Three independent experiments yielded similar results. (**d**) IFA of chlamydial OmcB expression. Cells were fixed at 40h pi, stained with anti-OmcB, followed by visualization of OmcB (red), P_*ompA*_-GFP (green), and DAPI-stained DNA (blue). Scale bar, 10 µm. Statistical significance was assessed by one-way ANOVA with Tukey’s post hoc test. ***P ≤ 0.001; ****P ≤ 0.0001.

To distinguish lethal effects from reversible arrest, cultures exposed to 1 µg/mL Mox from 16-48 hpi were transferred to drug-free medium and incubated for an additional 96 h. Under these conditions, the inclusion expansion resumed and EB production increased (Figure S1), indicating that a subset of organisms enters a reversible persistent state rather than undergoing irreversible damage. This outcome is consistent with previous observations using ofloxacin (5).

These results show that *Chlamydia* sensitivity to Mox is stage- and drug concentration-dependent and closely linked to DNA supercoiling dynamics. Early-stage organisms are highly vulnerable, while a subset of mid-cycle Ctr retain sufficient topological flexibility to adapt and survive under Mox stress by entry into a persistent state.

### Moxifloxacin inhibits chromosomal DNA replication during the exponential phase

To determine whether growth inhibition reflects impaired DNA replication, chlamydial genomic DNA (CtgDNA) accumulation was quantified using quantitative polymerase chain reaction (qPCR). Primer sets targeting loci proximal to the origin of replication (*oriC*) (e.g., *topA, ompA*) and the terminus (ter) (g*roEL1, gyrBA*) were used (Fig. 2a). This design allows assessment of replication activity and progression across the chromosome of Ctr.

**Figure 2.**
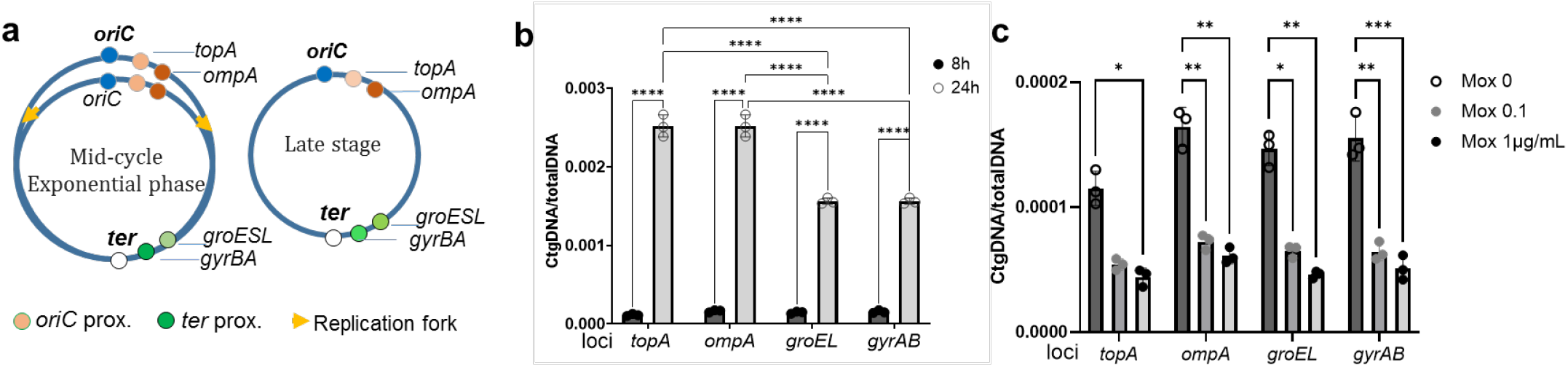
Effect of moxifloxacin on DNA replication. Total DNA extracted from Ct L2/434/Bu infected cells were used to quantify CtgDNA by qPCR. (**a**) Scheme of chromosomal loci proximal to the *oriC* and *ter* at exponential phase (mid-cycle) compared to late stage. (**b**) CtgDNA content increased from 8 to 24 hpi. (**c**) Mox reduced CtgDNA content from 8 to 24h. CtgDNA was normalized to total DNA from respective sample. Data represent mean ± SD from one representative experiment. Three independent experiments were performed with similar results. Statistical significance was determined using two-way ANOVA with the Tukey’s post hoc analysis; *P≤0.05, ***P ≤ 0.001, ****P ≤ 0.0001.

In unexposed cultures, CtgDNA increased steadily from 8 to 24 hpi, with enrichment of *oriC* proximal loci (Fig.2b). Exposure to Mox, either at 0.1 or 1 µg/mL, beginning at 8 hpi caused a reduction of CtgDNA at both *oriC* and *ter* (Fig.2c), indicating broadly reduced DNA replication. Similar inhibition occurred when Mox was applied from 16 to 24 hpi (Figure S2). Prolonged exposure (16–40 hpi) produced a greater CtgDNA decline, confirming sustained replication arrest.

These results show that DNA replication is most vulnerable during the exponential phase, when need for negative supercoiling is highest. Mox addition at this stage results in replication halt, underscoring the essential role of gyrase function and DNA topology in Ct replication.

### Topoisomerase gene expression responds rapidly and differentially to Mox exposure

The study of gene responses to supercoiling variations is often conducted by using a wild-type strain and inducing a rapid change in supercoiling with topoisomerase-inhibiting antibiotics (30, 33, 49). The same approach was employed to assess how Ctr topoisomerase regulation responds to Mox during mid-cycle. Wild-type Ctr L2/434/Bu cultures were exposed to increasing concentrations of Mox for 30 min (16-16.5h pi) (Fig. 3a), and transcript levels of *gyrAB* (encoding DNA gyrase), *parCE* (encoding Topo IV), and *topA* (encoding TopA) were quantified by RT-qPCR.

**Figure 3.**
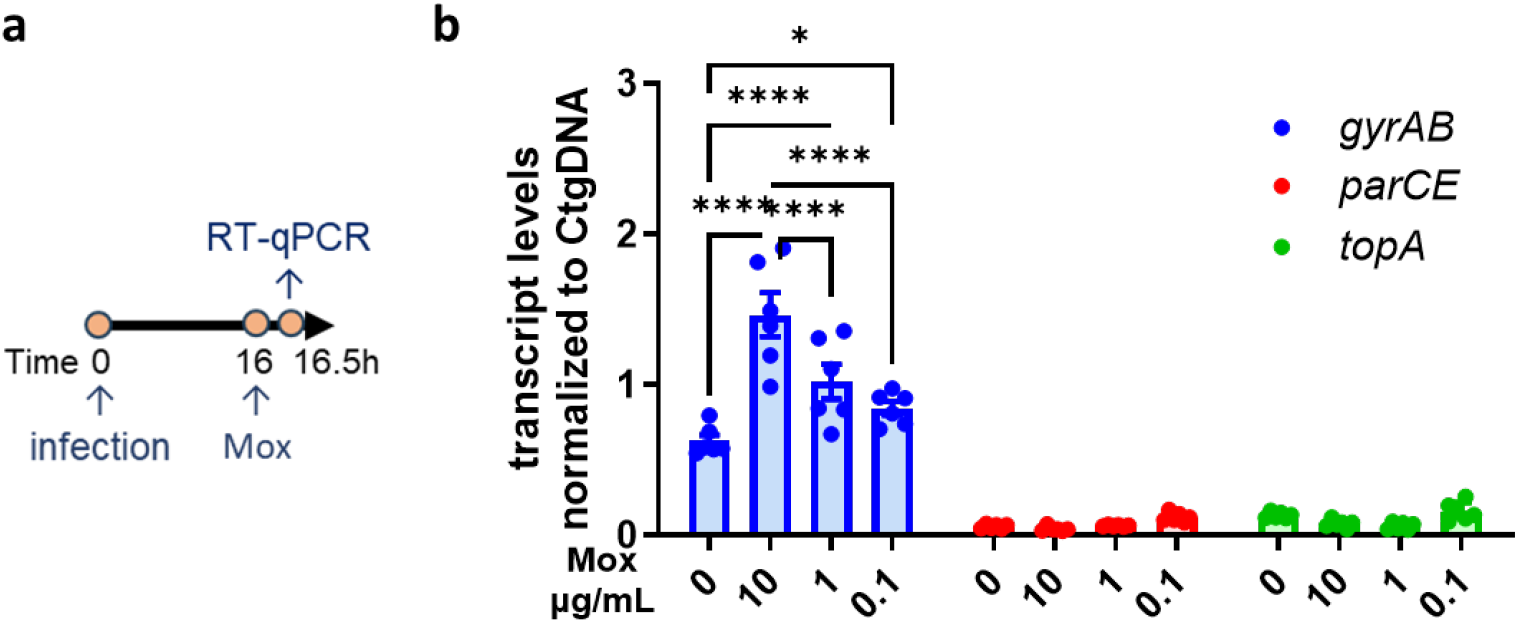
Moxifloxacin elicits transcriptional responses differently in topoisomerase genes. **(a)** Schematic diagram of the experimental procedure. Ctr L2/434/Bu cultures were exposed to Mox starting at 16 hpi to 16.5hpi. Unexposed cultures were used as controls. Total RNA was extracted, cDNA was made, and RT-qPCR analysis was performed on cDNA. (**b**) Changes in relative transcript levels of genes as indicated. Showings are results from Ctr cultures exposed to Mox (at concentrations of 10, 1, 0.1, 0 µg/mL). Quantified transcripts by RT-qPCR were normalized to the CtgDNA value from respective samples by qPCR with the same primer sets. Data represent means ± SEM from two individual experiments; each had technical replicate. Statistical significance was determined by two-way ANOVA. P⩽0.05, ***P ⩽ 0.001; ****P ⩽ 0.0001.

Mox caused a rapid, dose-dependent increase in *gyrAB* transcription (Fig. 3b), indicating stimulation of *gyrAB* expression in response to loss of negative supercoiling or DNA relaxing. By contrast, *topA* and *parCE* transcript levels showed little change. These results demonstrate selective induction of the gyrase during acute topological stress induced by Mox, suggesting that gyrase serves as the principal sensor and compensatory effector for restoring supercoiling balance.

### Moxifloxacin depletes GyrA protein independently of transcriptional changes

Because transcript-level changes alone could not explain the severe phenotypic defects, we assessed the abundance of GyrA protein, the catalytic subunit of DNA gyrase, by immunoblotting. To detect it, we generated a rabbit polyclonal anti-GyrA antibody targeting the C-terminal region (residues 818-836). Mock-infected HeLa cells served as negative controls to validate antibody specificity. Chlamydial Hsp60, a general stress chaperone, and host glyceraldehyde-3-phosphate dehydrogenase (GAPDH) were used as controls for bacterial and host protein, respectively.

Immunoblotting detected a ~94-kDa band corresponding to the expected size of GyrA (Fig. 4a–c). In unexposed cultures, GyrA levels increased from 16–24 hpi; in parallel, Hsp60 accumulated (Fig. 4a), correlating with increased needs for negative supercoiling and protein folding assistance. Mox exposure during this period caused a concentration-dependent depletion of GyrA, yet only modestly affected Hsp60 levels (Fig. 4b), indicating that Mox selectively depletes DNA gyrase despite *gyrAB* transcriptional induction following Mox exposure (Fig. 3).

**Figure 4.**
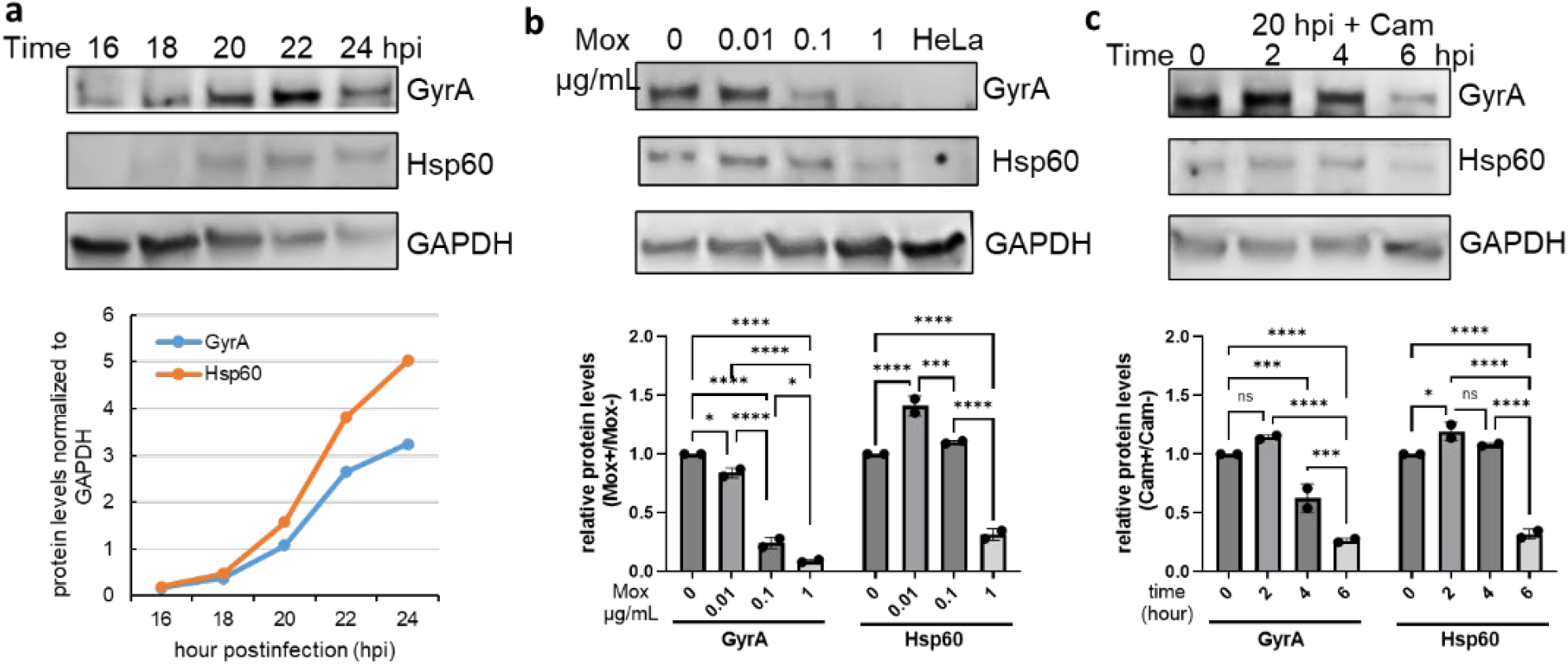
Moxifloxacin reduces GyrA levels in Ctr. Lysates from Ctr L2/434/Bu infected HeLa cells were analyzed by immunoblotting. Densitometry of protein bands were analyzed by ImageJ. (**a**) Time-dependent accumulation of GyrA and Hsp60 at indicated time points; each normalized to GAPDH. **(b)** Effects of 6 h Mox exposure (16–22 hpi) on GyrA and Hsp60 at the indicated drug concentrations. Relative protein levels were obtained by normalizing the value of GyrA/GAPDH or Hsp60/GAPDH from each sample to that in Mox-unexposed control (set to 1). Mock-infected HeLa cells were used as negative control. (**c**) Assessment of GyrA protein stability under chloramphenicol (60 µg/mL) starting at 20hpi for 2, 4, or 6 h. Ratios indicate fold changes relative to the Cam-untreated control (set to 1).

To determine whether the observed GyrA loss reflected increased degradation, we examined GyrA stability under chloramphenicol-mediated translation arrest. GyrA levels declined gradually over 2-6 hours (Fig. 4c) but far less than with Mox. The reduction in the amount of detectable GyrA protein may be indicative of the trapping of DNA gyrase in stable, Mox-bound cleavage complexes as reported previously in other bacteria (50, 51).

These protein level results demonstrate that Ctr DNA gyrase is a primary and sensitive target of Mox. The relative stability of Hsp60 in contrast to the low detectable GyrA protein suggests that rapid gyrase depletion may precede global proteome instability.

### GyrA inhibition is accompanied by TopA reduction

Because gyrase and TopA jointly maintain supercoiling homeostasis, we examined whether Mox-induced GyrA loss also affects TopA abundance. Endogenous TopA expression is most prominent during mid- to late-cycle (46). Immunoblotting confirmed detectable levels of TopA in Ctr between 22 and 26 hpi, although its abundance was relatively lower than that of GyrA (Fig. 5a). Exposure to Mox from 22 to 26 hpi led to a reduction of TopA levels, similar to the decreased levels of GyrA (Fig. 5b).

**Figure 5.**
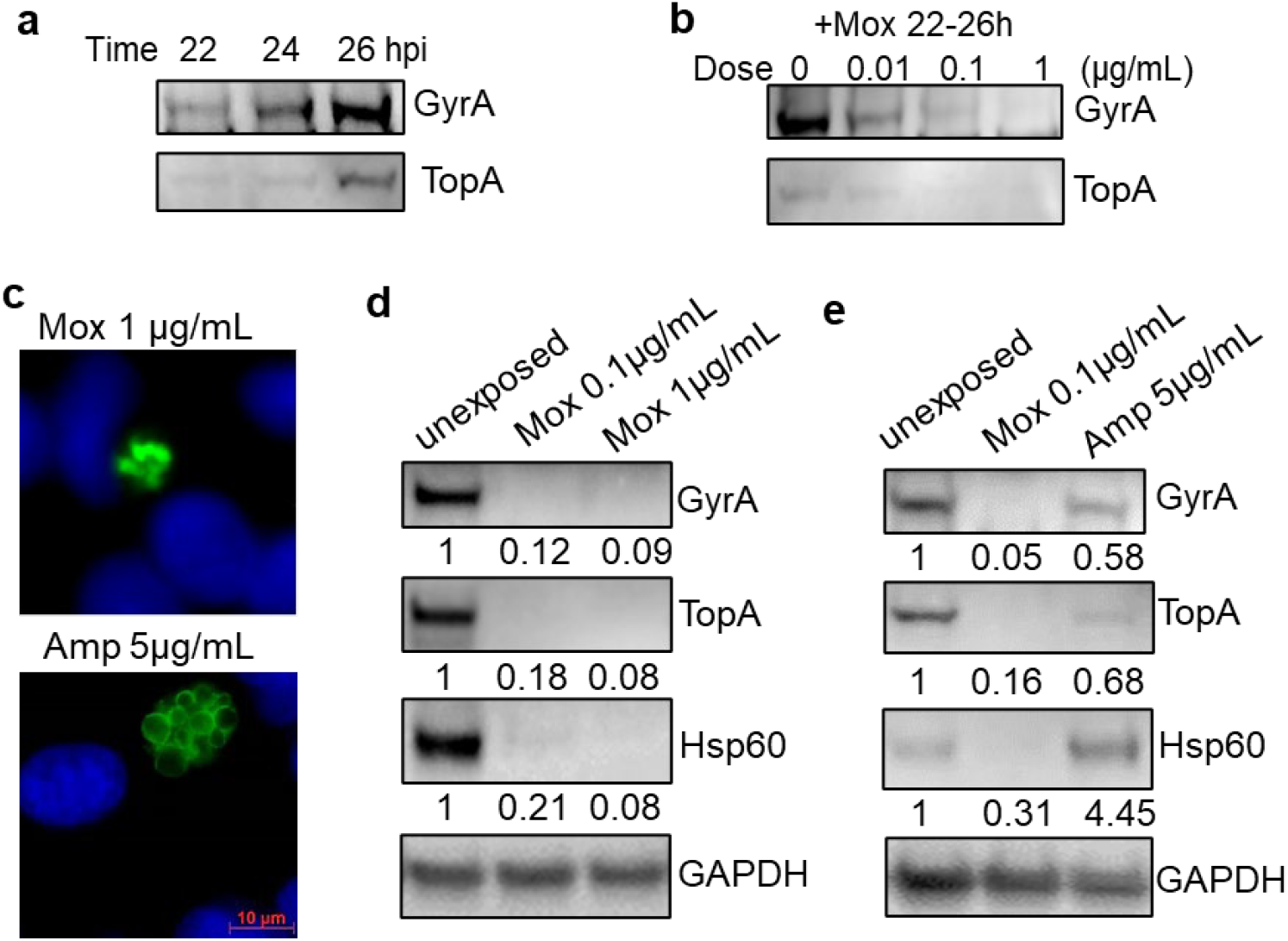
Reduction of TopA following Mox-induced GyrA inhibition. Lysates from wild-type Ctr strain L2/434/Bu infected HeLa cells were analyzed by immunoblotting to assess changes in TopA and GyrA. (**a**) Accumulation of GyrA and TopA in Ctr between 22 and 26 hpi. (**b**) Mox dose-dependent reductions of the TopA and GyrA levels from 22 to 26hpi. (**c**) Immunofluorescence image of Ctr exposed to Mox or ampicillin from 16 to 40 hpi. Fixed cells at 40h pi were immunostained to visualize the major outer membrane protein (MOMP; green) and DAPI-stained DNA (blue). Scale bar, 10 µm. (**d**) Immunoblot analysis of GyrA, TopA, and Hsp60 in Ctr exposed to 0.1 or 1 µg/mL of Mox at 40 hpi. Host GAPDH was used as loading control. (**c**) Comparing protein profiles under Mox- or ampicillin-induced stress. Densitometry of protein band was analyzed by ImageJ and then normalized to GAPDH from the same sample. Ratios represent fold changes relative to untreated controls (set to 1).

To assess the effects of prolonged Mox exposure and determine whether Ctr specifically responds to topological cues, cultures were treated with Mox from 16 to 40 hpi. As a control for envelope stress, cultures were exposed to ampicillin. Ampicillin induced enlarged but morphologically distinct persistent forms, accompanied by modest reductions in GyrA and TopA and a pronounced increase in Hsp60 (Fig. 5c–e). In contrast, Mox treatment led to concomitant decreases in GyrA and TopA at 40 hpi, along with reduced Hsp60 levels, suggesting overall proteome instability.

Therefore, the coordinated loss of gyrase and TopA appears to be a specific consequence of Mox-induced topological stress rather than a general stress response. These data further support that Mox fundamentally disrupts the enzymatic balance required to maintain supercoiling homeostasis.

### Supercoiling disruption reprograms promoter-specific transcription

Our findings thus far demonstrate that DNA supercoiling acts as a critical regulator of multiple chlamydial processes and that Mox disrupts such regulation. To determine how supercoiling disruption affects virulence-associated loci, we quantified transcript levels of the Hsp60 gene (*groEL1*), its co-transcribed partner *groES*, and the MOMP gene (*ompA*). Because of their established roles in pathogenesis and stress adaptation, these genes are well-characterized in models of stressor-induced chlamydial persistence (15, 52–54). To interpret the impact of Mox on early transcriptional dynamics, we initially evaluated the effects of exposure of wild-type Ctr L2/434/Bu to Mox (0.2 µg/mL) for either 0.5 h or 8 h, followed by RT-qPCR analysis.

As shown in Fig. 6a, thirty-minute exposure resulted in reduced transcript levels at both *ompA* and *groESL1* loci. However, the responses diverged significantly over time: after 8 h of exposure, *groESL1* transcript levels increased sharply, whereas *ompA* were unchanged or remained at low levels. These data indicate that topological stress triggers a gene-specific adaptation, with a preferential shift toward enhanced protein-folding capacity by upregulating *groESL1*.

**Figure 6.**
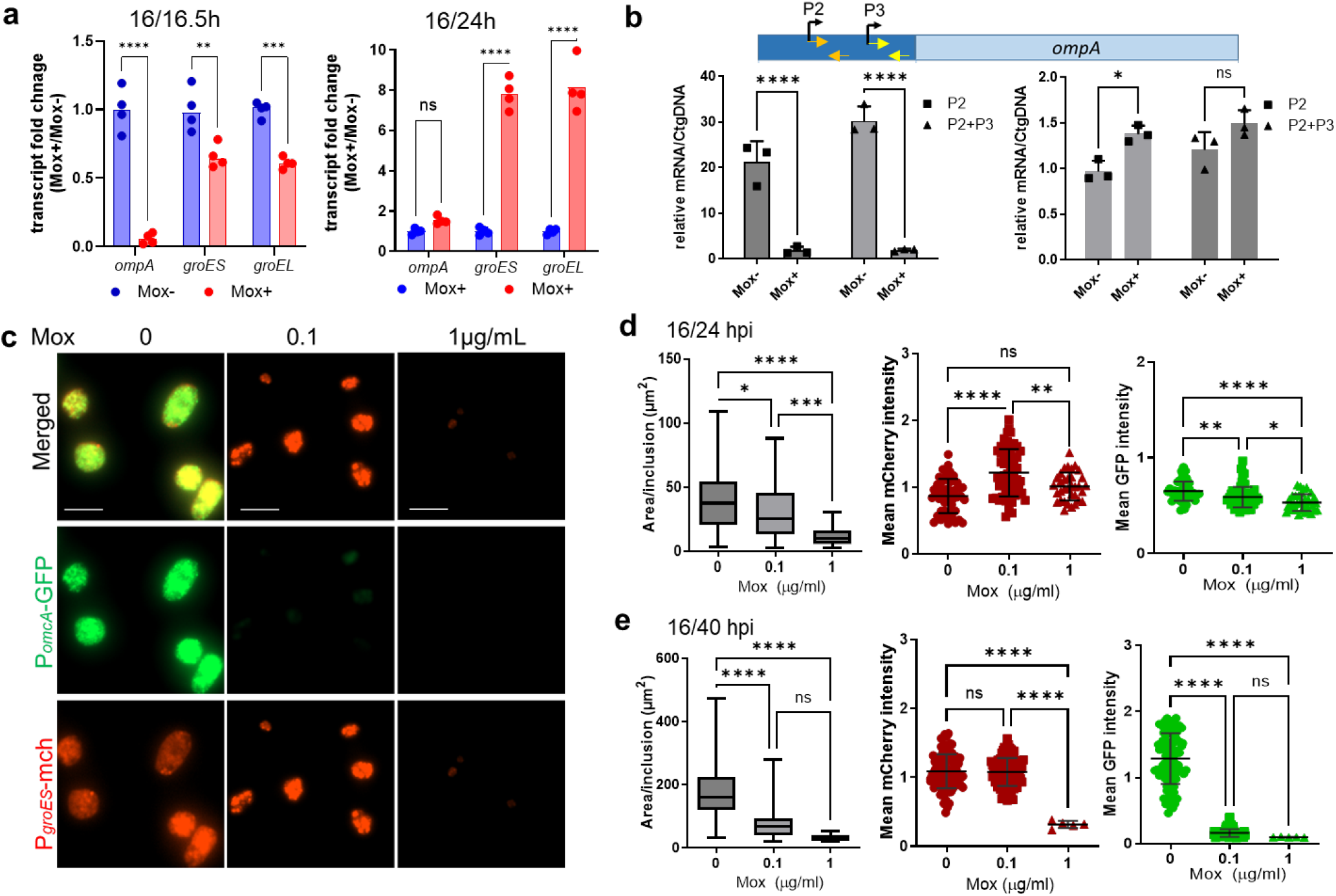
Mox impacts promoter-specific transcription in Ctr. **(a)** Fold changes in transcript levels of *ompA* and *groESL1* after 0.5 h or 8 h of exposure to Mox at 0.2 µg/mL compared with unexposed controls (set to 1). Data represent means ± SD from an independent experiment. Three independent experiments were performed with similar patterns. (**b**) Analysis of *ompA* P2- and P2+P3-derived transcripts after 0.5 h (left panel) or 8 h (right panel) Mox exposure. Orange or yellow arrows show the locations of primer sets used for amplifying the P2 or P2+P3 derived transcripts. **(c)** Representative live-cell fluorescence images of reporter strain exposed to increasing doses of Mox for 24 h (16–40 hpi). Scale bar, 10 µm. **(d, e)** Quantitation of Ctr inclusion area and mean mCherry or GFP fluorescence intensity (MFI) after exposure to Mox for 8 h (**d**) or 24 h (**e**). Data represent means ± SD from three biological replicates. Statistical significance was determined by one-way ANOVA with the Tukey’s post hoc test. *P≤ 0.05; **P≤ 0.01; ***P ≤ 0.001; ****P ≤ 0.0001; ns, not significant.

To investigate the basis for the *ompA* expression, we assessed the activities of its two σ^66^-dependent-promoters, P2 and P3 (48), using promoter specific primers (Table S2). Both P2 derived- and P2+P3-derived transcripts were reduced within 30 min of Mox exposure (Fig. 6b, left panel), mirroring the overall decrease in *ompA* transcription (Fig. 6a). At 8 h, a modest recovery was observed only from the P2 promoter (Fig. 6b, right panel), suggesting its potential role in maintaining minimal *ompA* transcription under Mox stress.

To determine whether the biphasic *groESL1* response reflects a transition from an immediate shock reaction to adaptive regulation, we monitored dynamic promoter activity together with Ctr growth patterns across increasing concentrations of Mox (Fig. 6c–e). For this analysis, we used a dual-reporter strain expressing mCherry under control of the *groES* promoter (P_*groES*_) and GFP under the *omcA* promoter (P_*omcA*_) (55). In untreated cells, mCherry fluorescence was detectable by 8 hpi, whereas GFP signal appeared by 16 hpi, consistent with the expected early and late promoter activities of P_*groES*_ and P_*omcA*_, respectively. Upon Mox exposure, P_*groES*_-driven mCherry levels remained stable or increased with rising Mox concentrations, consistent with the transcript analyses of *groESL1* in chromosomal context (Fig. 6a). In contrast, P_*omcA*_-driven GFP expression was strongly suppressed, corroborating the IFA observations (Fig. 1c). Only during prolonged and high-dose exposure (1 µg/mL to 40 hpi) did both reporters decline, indicating either global transcriptional inhibition or unstable proteome affecting the fluorescence reporters.

These results indicate that Mox-induced DNA supercoiling disruption produces gene-specific transcriptional outcomes rather than uniform repression and that DNA supercoiling plays a central role in coordinating transcriptional programs in Ctr.

## Discussion

This study demonstrates that DNA supercoiling is a dynamic regulatory parameter that shapes chlamydial physiology throughout its developmental cycle. Mox imposes stage-specific sensitivity, inhibits chromosomal DNA replication, depletes DNA gyrase, reduces TopA levels, and drives promoter-specific transcriptional changes in Ctr. Because bacterial viability and nucleoid architecture depend on a balanced interplay between gyrase-mediated negative supercoiling and TopA-driven relaxation (33–36), perturbing these enzymes disrupts supercoiling homeostasis. Our findings support a model (Fig. 7) in which topological balance acts as a key determinant of Ctr developmental progression. Disrupting this balance influences multiple processes in Ctr and promotes its entry into a persistent state as a potential survival mechanism.

**Figure 7.**
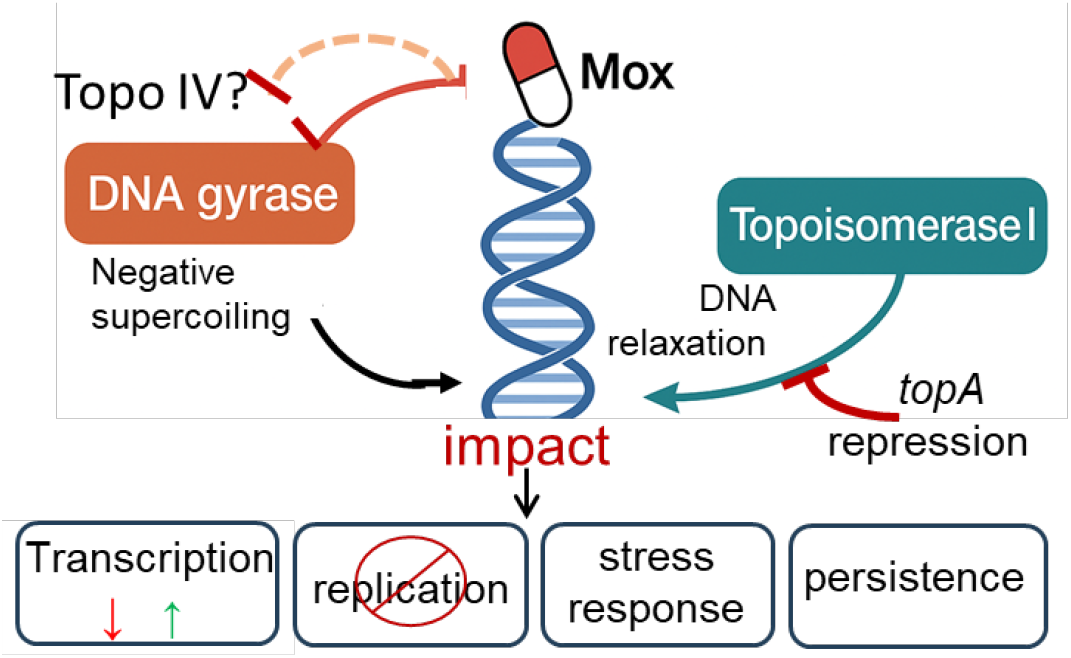
DNA supercoiling dynamics and fluoroquinolone response in Ctr. DNA gyrase introduces negative supercoils into the chromosome, whereas topoisomerase I (TopA) counterbalances this activity by relaxing supercoils. Mox inhibits DNA gyrase (and Topo IV), thereby reducing negative supercoiling and disrupting the topological homeostasis required for normal chlamydial development. Conversely, targeted repression of *topA* using CRISPRi perturbs DNA topology in the opposite direction by increasing negative supercoiling (44). These findings highlight that coordinated gyrase and TopA activities are central to maintaining supercoiling homeostasis in *C. trachomatis*. Disruption of this balance leads to growth arrest, formation of persistent forms, impaired DNA replication, and promoter-specific transcriptional shifts.

Prior work shows that DNA supercoiling is a global regulator of gene expression and developmental progression in *Chlamydia* and other bacteria (30, 31, 56, 57). Our findings extend this concept by showing that Ctr exhibits stage-specific susceptibility to Mox, reflecting changes in the topological levels associated with its developmental cycle. Early Mox exposure disrupts the rapid restoration of negative supercoiling needed as the EB chromosome decondenses to a transcriptionally active state during germination. Mid-cycle Ctr (mainly RBs) experiences large torsional strain due to high levels of replication and transcription. A subset of RBs may enter a persistent state rather than undergo lethal damage, indicating the sufficient capacity to preserve chromosome integrity despite impaired gyrase activity induced by Mox. By late stages, reduced drug sensitivity coincides with chromosome re-condensation and diminished reliance on gyrase-mediated supercoiling. Together with our CRISPRi-mediated *topA* knockdown data (47), these results support a model in which altered supercoiling modulates fluoroquinolone sensitivity and governs transitions between vulnerable, replicative, and persistent states.

Extending this stage-specific perspective, the essential role of DNA topology in Ctr chromosome replication becomes evident from the strong inhibition of gDNA accumulation upon Mox exposure (Fig. 2a–c). Progressing replication forks generate positive supercoils ahead of the fork that must be resolved by gyrase-mediated negative supercoiling. Inhibition of gyrase disrupts this balance, increasing torsional strain, stalling replication forks, and decreasing genomic copy number in Ctr. The marked replication inhibition during exponential phase aligns with the high topological needs of active replication and underscores the dependence of chlamydial DNA replication on proper DNA supercoiling levels.

In parallel, topological stress elicited a transcriptional response of topoisomerase genes (Fig. 3), highlighting the intimate link between DNA topology and its enzymatic regulators. In model organisms such as *Escherichia coli*, gyrase expression operates within a well-characterized negative feedback loop in which DNA relaxation stimulates transcription of gyrase genes to restore supercoiling homeostasis (58, 59). This self-limiting regulatory circuit ensures that gyrase levels rise only transiently, sufficient to counteract DNA relaxation. *Chlamydia*, however, appears to employ a fundamentally different regulatory logic. Its topoisomerases are expressed in a temporally coordinated manner across the developmental cycle. Previous *in vitro* transcription assays suggest an unconventional positive feedback mechanism, whereby Ctr gyrase gene promoters are more active on supercoiled DNA templates (30). Our Ctr study shows that Mox exposure elicited a dose-dependent induction of *gyrAB* transcription, indicating that excessive DNA relaxation can override the activation threshold required to trigger transcription from gyrase promoters, consistent with the canonical negative feedback model described in *E. coli*. In contrast, transcript levels of *topA* and *parEC* remained unchanged, suggesting that under conditions of acute topological stress, Ctr preferentially prioritizes restoration of negative supercoiling primarily through gyrase activity rather than engaging other topoisomerase functions.

Protein-level measurements further reinforce DNA gyrase as the primary functional target of Mox (Figs. 4,-5). Fluoroquinolones kill bacteria by trapping DNA gyrase (and Topo IV) in drug-enzyme-DNA cleavage complexes (40, 50, 51), which inhibits DNA replication and halts bacterial growth. Because the enzymes (including the GyrA) are trapped on the chromosome, the detectable enzyme level is reduced. Moreover, the genomic instability caused by severe topological stress may broadly decrease the stability of the entire proteome. These findings complement earlier results (Fig. 3) demonstrate that depletion of GyrA can dominate Ctr outcomes even when *gyrAB* transcripts are sustained or induced. Although a lack of antibodies prevented direct measurement of GyrB levels, coordinated loss is possible, given the co-transcribed nature of *gyrA* and *gyrB* and their obligatory function as a DNA gyrase holoenzyme (GyrA_2_GyrB_2_).

Notably, simultaneous decreases in GyrA and TopA protein suggest coordinated tuning of negative and positive supercoiling activities, a protective mechanism to prevent excessive relaxation when gyrase function is impaired (33–36). The absence of a similar response under ampicillin-induced envelope stress confirms that Ctr is able to sense and distinguish topological stress from other stressors and tailor its topoisomerase regulation accordingly.

While both gyrase and Topo IV enzymes can be fluoroquinolone targets, the dominant target varies by organism (37, 40): DNA gyrase in Gram-negative species (e.g., *E. coli*) and Topo IV in Gram-positive species (e.g., *Staphylococcus aureus*). Although Topo IV genes exhibited transcriptional behaviors distinct from those of gyrase, our data do not exclude Topo IV as a relevant Mox target in Ctr, particularly under conditions that promote DNA cleavage-complex formation. Clarifying Topo IV’s contribution to the chlamydial developmental cycle remains an important direction for future work.

Beyond topoisomerases, our results demonstrate that even modest perturbations in supercoiling generate rapid and divergent transcriptional responses in Ctr, exemplified by the contrasting behaviors of *ompA* and *groESL1* (Fig. 6a-e). These data indicate that Ctr relies heavily on topological cues to coordinate developmental progression and to initiate stress-adaptive programs. The strong induction of *groESL1* contrasts with the repression of *ompA* and *omcB*, suggesting that promoter architecture encodes gene-specific sensitivity to varying superhelical density. Promoter-focused analyses further support this conclusion by showing that topological stress does not uniformly suppress transcription. Instead, promoter structure and genomic context modulate the magnitude and direction of transcriptional responses, enabling a rapid shift from growth-associated to stress-responsive gene expression. Given Ctr’s streamlined genome, limited repertoire of transcription factors, absence of canonical stress sigma factors, and broad diversity of promoter architectures (21), promoter-encoded topological sensitivity may constitute an important regulatory axis. Such a mechanism would allow Ctr to adjust transcription swiftly and efficiently in response to environmental conditions, developmental progression, and antibiotic-induced supercoiling stress.

Although this study focuses on Mox, other fluoroquinolones induce similar effects on Ctr (5, 45),, underscoring the broader relevance of topological stress as a unifying mechanism driving developmental disruption. Future work aimed to profile global transcriptional shifts, map genome-wide supercoiling states, identify topologically sensitive promoters, and elucidate how topoisomerases interface with developmental regulators will be critical for advancing our understanding of Ctr pathobiology. Such efforts may also reveal opportunities to exploit vulnerable developmental windows or to intervene in persistence-associated pathways, thereby informing the development of improved therapeutic strategies.

## MATERIALS AND METHODS

### Chemicals

Antibiotics and chemicals were purchased from MilliporeSigma (St. Louis, MO, USA). Primers were synthesized by Integrated DNA Technologies (Coralville, IA, USA).

### Cell cultures and bacteria growth

HeLa 229 cells (human cervical epithelial carcinoma cells; ATCC CCL-2) were cultured in Dulbecco’s Modified Eagle Medium (DMEM) supplemented with 1 mM glutamic acid, 5% fetal bovine serum and 20 μg/ml gentamycin (DMEM-5) at 37 °C in an incubator supplied with 5% CO2. Cells were confirmed to be Mycoplasma negative by PCR as described previously (60).

Ctr strains used are: wild-type L2/434/Bu, L2/P_*ompA*_-GFP (single reporter)(48), and the L2/p2TK2-SW2 mCh(GroL2)GFP(OmcAL2) (dual reporter) (55) (Table S1). When approximated, ampicillin (5 µg/mL) and spectinomycin (500 µg/mL) were used for selection of transformed strains. Propagation and purification of WT and transformed *Chlamydia* organisms were performed as described previously (61). The purified EB pellet was resuspended in sucrose-phosphate-glutamic acid buffer (10 mM sodium phosphate, 220 mM sucrose, and 0.50 mM L-glutamic acid). The EB aliquots were stored at −80°C until use.

For growth phenotypic analysis, EBs were used to infect cells in 96-well plates (catalog #655090, Greiner) with a dose that results in ~30% of cells being infected. After centrifugation with a Beckman Coulter model Allegra X-12R centrifuge at 1,600 × g for 45 min at 37°C, fresh medium was added to the infected cells and incubated at 37°C for various time periods as indicated in each experimental result. To generate persistence model, Ctr infected HeLa cells were cultured in medium containing ampicillin at 5 µg/mL to induce envelope stress. Mox was used to produce topological stress. Mox stock solution was made in 100% DMSO at 10 mg/mL and diluted in the corresponding culture medium at the concertation for use. The control medium lacking Mox contains an equal percentage of DMSO.

### Antibodies

A rabbit polyclonal anti-GyrA serum was produced with synthesized polypeptide corresponding to residues 818–836 at the C-terminal region of GyrA (Pacific Immunology, Ramona, CA). The final serum was purified by an affinity column. The aliquots with 40% glycerol were stored at −20°C prior to use. The other primary antibodies used are (i) a mouse monoclonal anti-MOMP antibody (L2I-45); (ii) a mouse monoclonal anti-Hsp60 antibody; (iii) rabbit polyclonal anti-OmcB (46); (iii) a mouse polyclonal ant-TopA antibody (46); and (iv) a mouse monoclonal antibody to GAPDH (catalog #T5168, MilliporeSigma). The secondary antibodies used are (i) Alexa Fluor 568-conjugated goat anti-mouse IgG (catalog #A11004) from Invitrogen (Carlsbad, CA, USA) and (ii) horseradish peroxidase (HRP)-conjugated goat anti-rabbit IgG (catalog #213204, Abcam) and HRP-conjugated anti-mouse IgG (catalog A0168, MilliporeSigma).

### Microscopy

Automated live-cell images in 96-well culture plates were acquired using an imaging reader Cytation1 (BioTek Instrument). Gen5 software was used to process and analyze the inclusion morphology (e.g., inclusion size, numbers, and mean fluorescence intensity [MFI]).

For indirect IFA, HeLa cell monolayers grown in a glass culture chamber (NUNC) or a coverslip in 24-well culture plate were infected with *C. trachomatis* to achieve ~30% infection. At various times post infection, the cultures were fixed with 4% (wt/vol) paraformaldehyde dissolved in phosphate-buffered saline (PBS) (pH 7.4) for 15 min at room temperature, permeabilized with 0.1% (vol/vol) Triton X-100 for an additional 15 min, and blocked with 2% (wt/vol) bovine serum albumin in PBS for 30 min. Then, cells were incubated with the indicated primary antibody overnight at 4°C, followed by incubation with Alexa Fluor 488/568-conjugated secondary antibody for 45 min at 37°C. DAPI was used to label DNA. In some experiments, cell images were photographed using an inverted fluorescence microscope (Zeiss Axio Observer D1) and analyzed with AxioVision software, version 4.8.

### Inclusion forming unit assay

To determine the infectious EB number in culture, samples were harvested by scraping three wells of a 96-well plate in medium, and lysed via a single freeze-thaw cycle, serially diluted, and used to infect a fresh HeLa cell monolayer in 96 well plate. After 24 hr, cells were fixed with 4% paraformaldehyde, and stained with a mouse anti-MOMP antibody, followed by staining with Alexa Fluor 568-conjugated goat anti-mouse IgG. The inclusions were counted using Cytation 1. All experiments were performed three times for three biological replicates. The titers of recoverable EBs in cells were converted to inclusion-forming units per milliliter.

### Immunoblotting analysis

Ct-infected cells in 12-well culture plates were lysed directly in 8 M urea buffer containing 10 mM Tris-HCl (pH 8.0), 0.1% SDS, and 2.5% β-mercaptoethanol. The protein content was determined by a bicinchoninic acid protein assay kit (Thermo Fisher). An equal amount of protein was loaded into a single lane of the 4%–15% SDS-polyacrylamide gel (BioRad). After separation, proteins were transferred onto a PVDF membrane (Millipore) for immunoblotting with primary antibody, followed by incubation with horseradish peroxidase-conjugated secondary antibody. The protein bands were visualized by an enhanced chemiluminescence kit (Kindle Biosciences). The blot was imaged on an Azure c600 imaging system.

### Nucleic acid preparation and analysis

*C. trachomatis*-infected HeLa cells in 24-well plates were harvested and used to isolate DNA and RNA using quick DNA/RNA miniprep kit (catalog # D7001, Zymo Research). Residual DNA in the RNA samples was digested with 20-U RNase-free DNase I in-column for 30 min at room temperature, followed by extensive washing. A total of 2 µg of RNA per sample was reverse transcribed into cDNA using the high-capacity cDNA reverse transcriptase kit (catalog # 4368814, Applied Biosystems). The Fast SYBR green master mix (Applied Biosystems) was used for qPCR assay in 10 µL of reaction mixture on a real-time PCR system (Bio-Rad). Primer pairs used in this study were listed in Table S2. Each sample was analyzed in triplicate or quadruplicate in a 96-well plate. A negative control containing no Ct*g*DNA was included. The PCR cycle conditions were as follows: 50°C for 2 min, 95°C for 5 min, 95°C for 3 seconds, and 60°C for 30 seconds. The last two steps were repeated for 40 cycles with fluorescence levels detected at the end of each cycle. A standard curve was made from purified wild-type *C. trachomatis* L2/434/Bu genomic DNA with serial dilutions for each gene-specific primer pair. Relative transcripts per genome copy were then calculated as the number of transcripts divided by the number of chlamydial genome DNA contents measured by qPCR with the same primer set.

### Flow cytometry

HeLa cells were detached by trypsinization and fixed with 2% paraformaldehyde at room temperature for 20 minutes. Cells were then detected in a FACSVantage flow cytometer (BD Biosciences) using the FL-1 (green) channel. Flow cytometry data was recorded for at least 3 × 10^4^ cells per sample. Mock infection cells were used as a blank. Data were analyzed using FlowJo software Version 7.6 (TreeStar Inc) for both the percentage of GFP-expressing cell populations and mean fluorescence intensity.

### Statistical analyses

Data analyses were performed using GraphPad PRISM software as described in each result. Values of *P < 0.05 were considered statistically significant.

## Acknowledgments

We are grateful to Dr. Isabelle Derré (University of Virginia) for kindly providing reagents. C.D. is a scholar of NIH Postbaccalaureate Research Education Program (PREP) (1R25GM148309). This work is supported by a National Institutes of Allergy and Infectious Diseases grant R21AI175651 to L.S. and S.P.O.

